# Pathogen Detection and Microbiome Analysis of Infected Wheat Using a Portable DNA Sequencer

**DOI:** 10.1101/429878

**Authors:** Yiheng Hu, Gamran S. Green, Andrew W. Milgate, Eric A. Stone, John P. Rathjen, Benjamin Schwessinger

**Affiliations:** Research School of Biology, The Australian National University Acton ACT 2601, Australia; Wagga Wagga Agricultural Institute, New South Wales, Department of Primary Industry Wagga Wagga NSW 2650, Australia; ANU-CSIRO Centre for Genomics, Metabolomics and Bioinformatics, Acton ACT 2601, Australia

## Abstract

Fungal diseases of plants are responsible for major losses in agriculture, highlighting the need for rapid and accurate identification of plant pathogens. Disease outcomes are often defined not only by the main pathogen but are influenced by diverse microbial communities known as the microbiome at sites of infection. Here we present the first use of whole genome shot-gun sequencing with a portable DNA sequencing device as a method for the detection of fungal pathogens from wheat *(Triticum aestivum)* in a standard molecular biology laboratory. The data revealed that our method is robust and applicable to the diagnosis of fungal diseases including wheat stripe rust (caused by *Puccinia striiformis* f. sp. *tritici),* septoria tritici blotch (caused by *Zymoseptoria tritici)* and yellow leaf spot (caused by *Pyrenophora tritici repentis).* We also identified the bacterial genus *Pseudomonas* co-present with *Puccinia* and *Zymoseptoria* but not *Pyrenophora* infections. One limitation of the method is the over-representation of redundant wheat genome sequences from samples. This could be addressed by long-range amplicon-based sequencing approaches in future studies, which specifically target non-host organisms. Our work outlines a new approach for detection of a broad range of plant pathogens and associated microbes using a portable sequencer in a standard laboratory, providing the basis for future development of an on-site disease monitoring system.

---

Plant diseases, especially those caused by fungal pathogens, jeopardise global crop biosecurity. Increased global trade, human migration, environmental changes and the accelerated emergence of virulence have been identified as causes for increasing prevalence of diseases in plants and animals (Fisher et al. 2012). To prevent and manage diseases, rapid detection and identification of their causal agents are crucial. This is particularly important for outbreak management during the incursion of virulent new isolates that overcome classical methods of control such as the use of resistant crop varieties (Bhattacharya 2017; Pretorius et al. 2000). Traditional methods for crop disease diagnosis rely largely on the expertise of pathologists whom identify diseases initially by eye in the field. While molecular methods of pathogen detection also exist, these are generally only capable of identifying specific diseases with low amounts of variation (De Boer and López 2011). For example, the enzyme-linked immunosorbent assay (ELISA) have been applied for the detection of tree pathogen *Xylella fastidiosa* (Sherald and Lei 1991). Recently, other optical sensor-based methods have been developed for the detection, identification, and quantification of plant diseases (Mahlein 2015). This method has been used to help smallholder farmers for the disease and pest control (PlantVillage, https://plantvillage.psu.edu/). Conventional pathology, however, remains the primary means of pathogen identification and harbors certain limitations: 1. It relies heavily on the physical appearance of disease symptoms; 2. It has difficulty detecting pathogens that do not infect aerial parts of the plant; 3. Co-infections may lead to conflicting visual symptoms; 4. No method currently exists for the rapid identification of unknown pathogens during outbreaks -a limitation epitomized by the recent, devastating incursion of South American races of wheat blast fungus in Bangladesh (Callaway 2016).

DNA sequencing-based methods have been applied to address limitations in detection and monitoring of environmental microbial communities (Nezhad 2014). Such assays rely largely on amplification and sequencing of small conserved regions called meta barcodes from pathogen genomes, including the internal transcribed spacer region or *Elongation Factor 1 alpha (EF1α)* and *Beta-tubulin* genes for fungal species and 16S ribosomal RNA sequences for bacteria (Janda and Abbott 2007; Raja et al. 2017). Combining multiple conserved gene loci, known as multilocus sequence typing (Maiden et al. 1998), can discriminate closely related species based on single nucleotide polymorphisms (SNPs) within amplicon sequences. This has been used, for example, to distinguish different closely related species of *Candida* that pose distinct risks and epidemiological outcomes (Odds and Jacobsen 2008). Amplicon sequencing also allows for characterisation of the microbial community - the microbiome - within a sample. The microbiome can impact plant function, for example, on altering flowering time (Panke-Buisse et al. 2015) or promoting colonization of fungal pathogens such as *Botrytis cinerea* (Ritpitakphong et al. 2016). Furthermore, the microbiome is likely to contain organisms that interact with disease-causing agents and, as some of these organisms have anti-microbial activities against plant pathogens, could potentially be harnessed for biocontrol strategies (Ellis 2017). Restrictive read lengths generated by the lllumina sequencing platform limits sequence-based identification of microbial species. Standard Illumina sequencing only allows for a maximum amplicon length of 500 base pairs (bp) to be sequenced and, as such, only the genetic variation present within 500 bp is available for taxonomic assignment of the organism in question (Benítez-Páez and Sanz 2017). Use of PCR also biases the abundance of individual amplicons, making estimation of population frequencies problematic (Kennedy et al. 2014).

Next-generation DNA sequencing platforms have developed rapidly during the last half-decade, overcoming the read length issue and facilitating the identification of microbial species (Benítez-Páez and Sanz 2017). Multiple single molecule, long-read sequencing technologies are currently in use. Most predominant among these are single molecular real-time (SMRT) sequencing (PacBio) and nanopore sequencing (MinION; Oxford Nanopore Technologies (ONT)). PacBio SMRT sequencing relies on the detection of fluorescent nucleotides incorporated into a single DNA molecule during its synthesis within a nanoscale observation chamber (Ardui et al. 2018). Nanopore sequencing enables direct sequencing of native DNA by measuring voltage changes across an artificial membrane when a single DNA molecule passes through a nanopore embedded in the membrane of a flowcell (Jain et al. 2016). Advantages of nanopore sequencing include its relatively low cost, pocketsize form and the potential for real-time data analysis, while current disadvantages include relatively low per read accuracy of around 90% (Leggett and Clark 2017). Because the MinION is a mobile technology independent of large sequencing facilities, it has been useful for on-site sample sequencing in extreme environments such as microbial paleo mats in the Antarctic (Johnson et al. 2017). For rapid pathogen identification, studies have shown a turnaround time of six hours for bacterial from clinical samples with MinION (Charalampous et al. 2018), and from seconds to up to four hours for detecting cassava virus in Africa (Boykin et al. 2018) - both unachievable with previous technologies. For antibiotic resistance, Břinda et al. (2018) reported their identification of known resistant bacterial strain within five minutes from real time MinION data. Moreover, the MinION provides significantly longer sequencing reads (~10 kb) than other sequencing platforms such as Illumina, allowing for more genetic information per read and thus more precise identification of the organism from which the read is derived (Benítez-Páez, Portune, and Sanz 2016; Benítez-Páez and Sanz 2017). Although recent studies have shown important advances in identification of bacterial pathogens using nanopore sequencing from clinical samples (Mitsuhashi et al. 2017; Schmidt et al. 2017), little has been done testing this new sequencing platform in terms of plant pathogens detection and microbiome profiling of diseased plants.

Here, we tested nanopore sequencing for disease diagnosis in field-grown wheat plants and explored its potential for microbiome profiling. We designed a whole-genome shot-gun sequencing workflow for identification of major pathogen species from field samples and profiling of microbes present in these samples to the species level. As a proof-of-concept, we applied our sequencing workflow to wheat leaves infected with known fungal pathogens sampled from the field and a fungicide treated control. By mapping sequencing reads to databases, we successfully detected all fungal diseases present in the infected samples including stripe rust (caused by *Puccinia striiformis* f. sp. *tritici),* septoria tritici blotch (caused by *Zymoseptoria tritici)* and yellow leaf spot (caused by *Pyrenophora tritici repentis),* and found one disease (septoria nodorum blotch, caused by *Parastagonospora nodorum).* We also conducted preliminary characterisation of the microbiomes associated with those diseases, identifying the bacterial genus *Pseudomonas* as co-present with *Puccinia* and *Zymoseptoria* infections. Our results suggest that portable nanopore sequencing has a considerable potential for adaptation to a broad range of crop disease diagnoses and environmental monitoring applications under field conditions.

## MATERIALS AND METHODS

### Sample collection

Wheat cultivar Crusader (Advanta Seeds Pty. Ltd.) seeds were planted in the disease nursery field on May 2016 at the Wagga Wagga Agricultural Institute (New South Wales (NSW), 35°02’24.3"S 147°19’09.5"E) with daily irrigation. Wheat straw infected the previous season with septoria tritici blotch and yellow leaf spot was allowed to stand in the field to promote development of pseudothecia. This infected material was used to manually inoculate plants in the nursery field at the tillering growth stage (4-5 weeks post emergence) while stripe rust infections occurred naturally at jointing stage (7-9 weeks post emergence). Healthy or diseased plants were diagnosed by a plant pathologist from the NSW Department of Primary Industries via symptomatology before samples were collected during September 2016. Five different treatment groups were collected into five separate containers. These included four infection types: three single infections *(Puccinia striiformis* f. sp. *tritici, Z. tritici* and *Pyrenophora tritici repentis)* and one double infection *(Puccinia striiformis* f. sp. *tritici + Z. tritici),* and a fungicide-treated control. For each treatment group, three separate entire wheat tillers were pooled in one container and shipped to the Rathjen laboratory, Research School of Biology, The Australian National University. Each biological replicate (n=4) consisted of two independent leaf cuts (~100 mg) from each treatment group containing visible disease symptoms or not in case of the control. The two leaf of each biological replicate were transferred into 2 mL Eppendorf tubes containing one 5 mm metal bead. All tubes were labelled and stored at −80°C prior to DNA extraction.

## DNA extraction

DNA extractions were performed according to Hu (2016). Briefly, 100 mg leaf tissue was homogenized and cells were lysed using cetyl trimethylammonium bromide (CTAB, Sigma-Aldrich Cas. No: 57-09-0) buffer (added RNAse T1, Thermo Fisher, Cat#EN0541, 1000 units per 1750 μL), followed by a phenol:chloroform:isoamyl alcohol (25:24:1, Sigma-Aldrich, Cat#P2069) extraction to remove protein and lipids. The DNA was precipitated with 700 μL isopropanol, washed with 1 mL 70 % ethanol, dried for 5 minutes at room temperature and resuspended in TE buffer containing 10 mM Tris and 1 mM EDTA at pH 8. DNA extractions were performed independently for each biological replicate (n=4), with the operator blind to the identity of each sample. After extraction, DNA was purified with one volume of Agencourt AMPure XP beads (Beckman Coulter, Inc. Cat#BECLA63880) according to the manufacturer’s protocol and stored at 4°C. Quality and average size of genomic DNA was visualized by gel electrophoresis with a 1% agarose gel for 1 hour at 100 volts. Representative gel images for replicate 3 and 4 are shown in Supplementary Fig. S1. DNA was quantified by NanoDrop and Qubit (Life Technologies) according to the manufacturer’s protocol. DNA quality and purity was evaluated as described previously (Schalamun et al. 2018).

### Library construction and sequencing

For each biological replicate, one PCR-based barcoded 1D sequencing libraries were constructed using 1D PCR barcoding workflow (SQK-LSK107 and EXP-PBC001, batch number EP01.10.0005) as previously described in Hu and Schwessinger (2018) with the omission of the DNA shearing step. Briefly, a dA-tailing reaction was performed on 700 ng genomic DNA from each sample using the NEBNext Ultra II End-repair/dA-tailing kit (New England Biolabs, Catalog #E7645). Repaired DNA was cleaned up using one volume of AMPure XP beads, washed on a magnetic rack using 70% ethanol, and eluted with 31 μL nuclease-free water. Thirty μL eluate was used for barcode adapter ligation via the NEB Blunt/TA Ligase Master Mix (New England Biolabs, Catalog #M0367S) and Barcode Adapter Mix (ONT 1D PCR barcoding kit EXP-PBC001), followed by another wash step on one volume of AMPure XP beads. Twenty ng of the adapter-ligated template in 2 μL of nuclease free water was added to each barcoding PCR reaction with 50 μL LongAmp Taq 2X Master Mix (New England Biolabs, Catalog #M0287S), 46 μL water and 2 μL PCR Barcodes (ONT 1D PCR barcoding kit EXP-PBC001). The PCR was performed as follows: denaturation at 95°C for 3 min, 15 cycles of 95°C/15 s; 62°C/15 s; 65°C/150 s, a final extension step at 65°C for 5 min. Barcoded DNA was purified using one volume of AMPure XP beads and pooled into 1 μg fractions in a total of 45 μL to which 5 μL of DNA CS control (ONT 1D ligation sequencing kit SQK-LSK107) was added. A further dA-tailing reaction was performed as above followed by purification on with one volume of AMPure XP beads. DNA was eluted in 31 μL nuclease free water. Adapter ligation was carried out using 30 μL end-prepped DNA, 20 μL Adapter Mix (ONT 1D ligation sequencing kit SQK-LSK107) and 50 μL NEB Blunt/TA Ligase Master Mix. The adapter-ligated DNA library was purified with one volume of AMPure XP beads using the Adapter Beads Binding buffer (ONT 1D ligation sequencing kit SQK-LSK107) for washing and eluted in 15 μL of Elution Buffer (ONT 1D ligation sequencing kit SQK-LSK107). Sequencing reactions were performed independently for each biological replicate on a MinION flowcell (R9.4, ONT) connected to a MK1B device (ONT) operated by the MinKNOW software (version 1.5.2 and version 1.6.11). Each flowcell was primed with 1 mL of priming buffer comprising 480 μL Running Buffer Fuel Mix (RBF, ONT) and 520 μL nuclease free water. Twelve μL of amplicon library was added to a loading mix including 35 μL RBF, 25.5 μL Library Loading beads (ONT library loading bead kit EXP-LLB001, batch number EB01.10.0012) and 2.5 μL water with a final volume of 75 μL and then added to the flowcell via the SpotON sample port. The "NC_48Hr_sequencing_FLO-MIN106_SQK-LSK107" protocol was executed through MinKNOW after loading the library. The sequencing run was restarted after 12 h and stopped after 48 h.

### Sequencing analysis

Raw fast5 files were processed via the Albacore 2.0.2 software (ONT) for basecalling, barcode de-multiplexing and quality filtering (Phred quality score of >7) following manufacturer’s recommendations. All reads that passed quality filtering for each barcode were treated in parallel as follows: firstly, NanoLyse (De Coster et al. 2018) was used to remove lambda phage control DNA sequences; secondly, barcode and adapter sequences were trimmed from the ends of reads using Porechop (Wick, Judd, and Holt 2018). To identify middle adapter sequences by Porechop, a 95% threshold was set, and reads with middle adapter sequences were split into two sequences; thirdly, seqtk (https://github.com/lh3/seqtk) was used to convert the processed fastq file into fasta format for further analysis. The two nucleotide BLAST (blastn, Basic Local Alignment Search Tool, Altschul et al. 1990) searches were performed using the fasta files for each barcode as queries: first, against a custom genome reference database that contained the wheat reference genome (Clavijo et al. 2017), accessed 09/2017, and the genome sequence of three fungal species used in our study: *Puccinia striiformis* f. sp. *tritici:* Schwessinger et al. (2018), accessed 09/2017; Z *tritici:* Goodwin et al. (2011), version 2, accessed 09/2017; *Pyrenophora tritici-repentis:* Manning et al. (2013), unmasked assembly, accessed 09/2017. The genome of a fourth species suggested to be absent from the studied wheat growing region, *Parastagonospora nordorum,* as a negative control (Hane et al. (2007), accessed 09/2017. Individual reads were assigned to have originated from a specific DNA sequence based on their best blastn hit (e-value < 0.01) (Zhang et al. 2000). Reads that did not hit the custom-made database were captured using the filterbyname.sh script from the bbmap script package (https://sourceforge.net/proiects/bbmap/). These reads were used as queries to search against the National Center for Biotechnology Information (NCBI) nucleotide (nt) database (ftp://ftp.ncbi.nlm.nih.gov/blast/db/, downloaded 09/2017) using blastn and the same settings and species assignment criteria mentioned above. Two Python scripts (QC_and_BLAST.py and creating_final_dataframe.py) that combine all steps from quality control to BLAST outputs and construct dataframes containing statistical and information from BLAST searches for each read were deposited at https://github.com/Yiheng323/Pathogen <u>Detection and Microbiome scripts</u> to facilitate the reproducibility of our results. All basecalled fastq data for each sequencing run were deposited at NCBI Short Reads Archive (SRA) under accession number PRJNA493553. The sequencing summary files and final dataframes were deposited as .tab files for each replicate at https://doi.org/10.6084/m9.figshare.7138262. The python scripts used for generating Fig. 2-5 and Supplementary Fig. S2 - S7 were also deposited at https://github.com/Yiheng323/Pathogen <u>Detection and Microbiome scripts</u>.

**Fig. 1.**
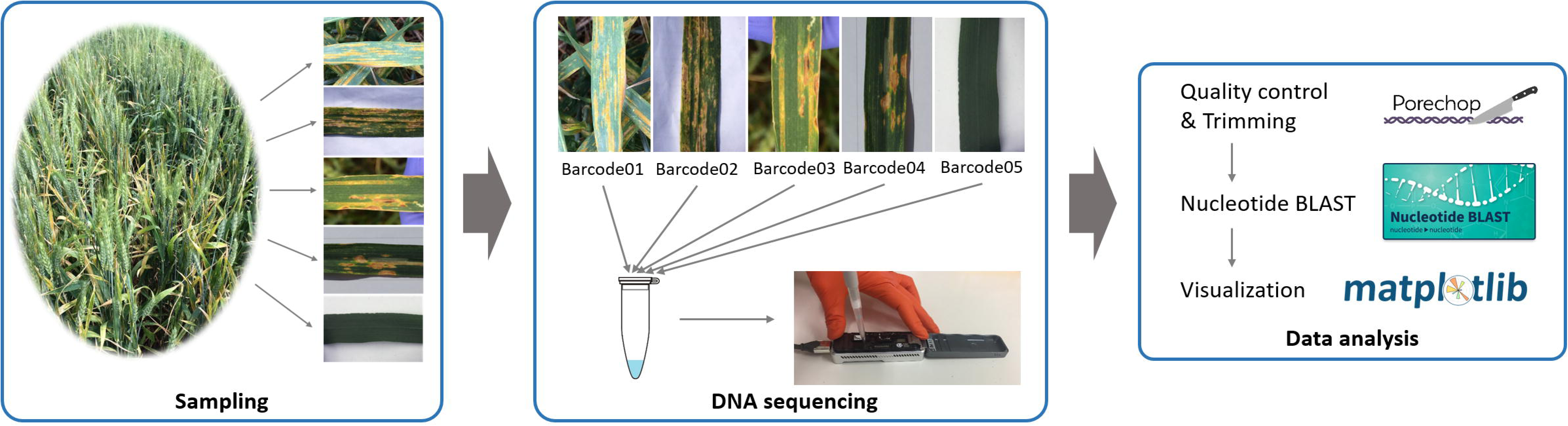
The Nanopore sequencing workflow for fungal pathogen identification. 1) Sampling: plant pathologists collected wheat leaf samples from disease nursery fields, with visual confirmation of disease symptoms for four infection types (three single fungal infections and one double fungal infection). Fungicide-treated wheat leaves of the same variety and growth stage were collected as a control treatment. 2) DNA sequencing: to minimise operator bias. DNA was extracted from all five samples without knowledge of the causative agent(s) affecting each sample. We labelled samples with DNA barcodes through a PCR reaction step before sample pooling and 1D library preparations (SQK-LSK107); 3) Data analysis: We quality-controlled sequences based on Phred quality scores and trimmed adapter sequence using Porechop (Wick, Judd, and Holt 2018). We performed BLAST (Altschul et al., 1990) searches using processed reads and identified the best hits. We summarized the total sequence length of hits to each species using the python *matplotlib* module. Images used in step 3 are from the official websites of each program.

**Fig. 2.**
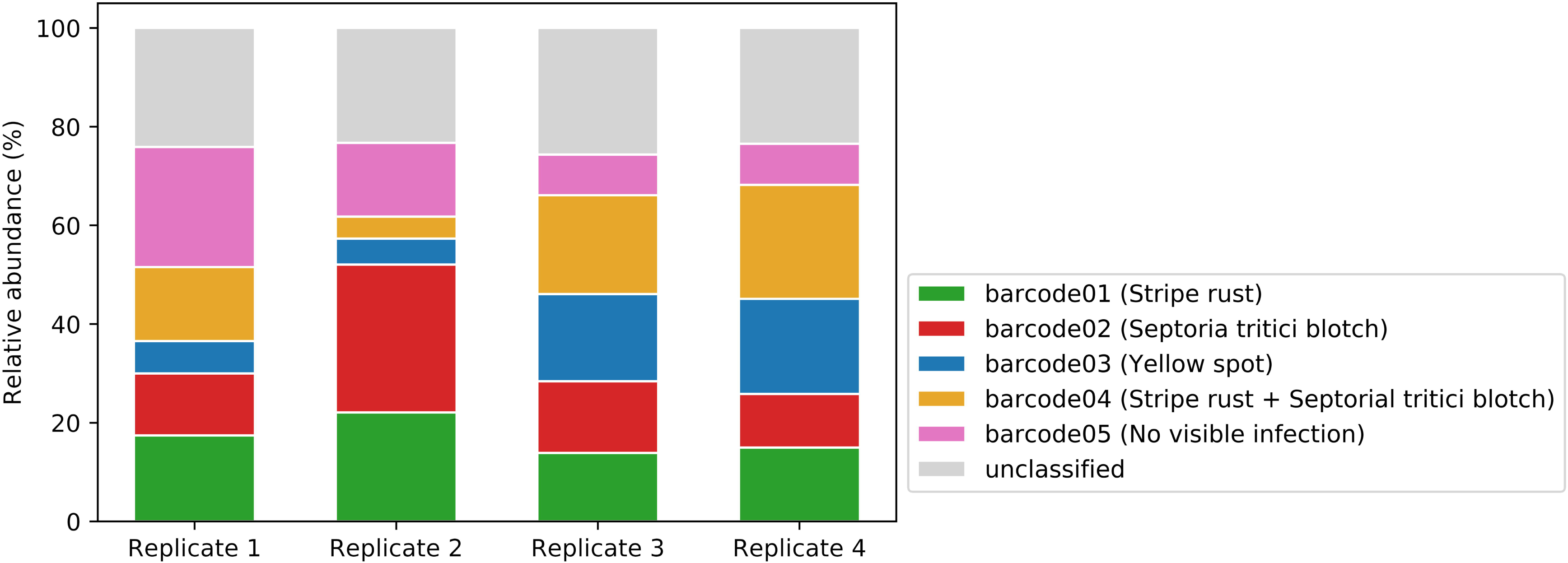
Uneven proportions of barcode-labelled DNAs across all biological replicates. Columns indicate four biological replicates from five different treatment types. The y-axis indicates relative abundance of total sequence lengths labelled with one of five DNA barcodes within each sample set. Each barcode represents one wheat sample. Reads without a detectable barcode were labelled as ‘unclassified’.

## RESULTS

### Sequencing showed variations in yield and consistency in barcode classification

We designed a pipeline for identification of three known wheat pathogens from infected field samples using MinION sequencing. We infected three experimental field plots as disease nurseries with their corresponding pathogens *(Puccinia. striiformis* f. sp. *tritici* for stripe rust disease, Z *tritici* for septoria tritici blotch and *Pyrenophora. tritici-repentis* for yellow leaf spot) commonly found in New South Wales, Australia, and conducted sampling during the 2016-17 growing season (September). The sequencing experiment included three steps (Fig. 1): 1. Pathologists collected three plants from each disease nursery field, with visual confirmation of disease symptoms from three single infections plus one double infection *(Puccinia striiformis* f. sp. *tritici + Z. tritici)* and a negative control comprising the same wheat variety treated with commercial fungicides at the 10- to 11-week growth stage (three months post germination). Individual plants were pooled by treatment type and shipped to the laboratory on ice for processing. 2. We extracted DNA from two independent leaf cuts of each treatment type without knowing the identity of each treatment to eliminate potential bias from operators. 3. We barcoded each sample to enable pooling of all five sample types into one sequencing run. The five wheat leaf samples were barcoded as follows: leaves infected with one of the three fungal pathogens (barcodes 1-3); leaves infected with two pathogens (barcode 4); and fungicide treated control without disease symptoms (barcode 5). We performed steps 2 to 3 four times independently as biological replicates, with Step 3 performed using a MinION 1D PCR barcoding workflow (EXP-PBC001 and SQK-LSK107). The raw data of each nanopore sequencing run were converted into fastq sequencing reads using Albacore software (ONT), with a quality (Q) score for each base estimating the likelihood of the identified base being correct using the Phred scale (Ewing et al. 1998; Ewing and Green 1998). Based on read mean Q scores, Albacore grouped reads into pass (Q score > 7) or fail (Q score <= 7) based on manufacturer’s recommendations. We binned reads based on the barcode sequences identified by Albacore at both ends of each sequencing read. If no barcode sequence could be identified, we named the corresponding read as unclassified. Overall, ~92-95% of the total sequence length of all four sequencing runs passed quality filtering (Supplementary Table 1). The total sequence length from the four runs varied from 682 to 5100 megabase pairs (Mbp). The mean read length ranged from 1380 to 3054 bp. Proportions of each barcode in each run were relatively even, although 23.3-25.7% of the sequences were unclassified in all four runs (Fig. 2).

### Sequence-based identification correctly identifies the fungal species causing wheat diseases

We performed blastn searches for all quality-filtered reads against a custom genome reference database. We assigned the origin of each read based on its best blastn hit within each database (e-value < 0.01) using a ‘winner-takes-all’ strategy which only considers the best match of each sequencing read. The custom genome reference database contained the wheat genome and four fungal genomes including *Puccinia striiformis* f. sp. *tritici, Z. tritici, Pyrenophora tritici-repentis* and *Parastagonospora nodorum.* These represent the host plant genome, three fungal species present in infected field samples, and one fungal species *(Parastagonospora nordorum)* not expected to be present in the sampling area as a negative control. The majority of reads were assigned to wheat sequences, comprising total sequence length of 90.1% in flowcell 1, 80.1% in flowcell 2, 92.7% in flowcell 3 and 91.4% in flowcell 4 (Fig. 3). We found that healthy leaf samples (barcode 5) contained no or minor total length percentages of pathogen-related sequences in all four replicates (< 0.01%, Fig. 4). This indicates that our analysis method is robust and does not lead to high background signals in healthy wheat samples (barcode 5). Significantly higher proportions of total sequencing length (0.5-5.7%) from infected samples (barcode 1-4) were assigned as pathogen genome-derived. In all cases, the pathogen genome identified by the highest total sequence length corresponded to the pathogen causing the disease as identified by symptomatology (Fig. 4). This indicates that we are clearly able to identify the disease-causing pathogen via our sequencing workflow in cases of single infections. Infections with two pathogens (barcode 4) were identified less clearly. In barcode 4 from all biological replicates, total sequence length percentages assigned to *Puccinia striiformis* f. sp. *tritici* were between 0.1% and 0.4%; close to or lower than the proportion assigned to negative control species *Parastagonospora nodorum.* The total relative sequence length identifying each major pathogen species within each barcode were similar across all replicates, with around 0.6% of barcode 1 from *Puccinia striiformis* f. sp. *tritici,* 1.9-5.0% of barcode 2 from Z *tritici,* 2.1-5.7% of barcode 3 from *Pyrenophora tritici-repentis,* and 0.1-0.4% of barcode 4 from *Puccinia striiformis* f. sp. *tritici* and 0.6-1.2% from Z *tritici.* Unexpectedly, we also identified *Parastagonospora nodorum* in samples derived from different treatment groups in all four biological replicates. To ascertain whether *Parastagonospora nodorum* was truly present, we extracted all 915 reads that were assigned to have originated from the *Parastagonospora nodorum* genome in all four biological replicates. We used these reads to perform an open-ended blastn search against the entire NCBI non-redundant nucleotide (nt) database. We found that 344 of 915 reads were assigned to genus *Parastagonospora,* including 339 reads identifing *Parastagonospora nodorum* SN15 genome (Supplementary Fig. S2). A further 371 reads hit related species, including 226 from the same *Pieosporaies* order: *Leptosphaeria* spp. (84 reads), *Aiternaria* spp. (81), *Pyrenophora* spp. (10), *Bipolaris* spp. (9), *Shiraia* spp. (9) and other genus (Supplementary Fig. S2). The remaining 200 reads fell above the given e-value threshold (0.01) and failed to be assigned. The difference between both BLAST searches potentially stems from the relatively high error rate of the MinION sequencer and the distinct conserved regions from genomes of closely related species (Jain et al. 2018; Laver et al. 2015). We tested for the presence of *Parastagonospora nodorum* in one of our four biological replicates using an independent molecular biology approach. We performed a PCR screen in replicate 3 using the *PnTox3* gene that is universally found in all Australian *Parastagonospora nodorum* isolates but not in any other fungal species (Liu et al. 2009). We were able to amplify a specific band corresponding to *PnTox3* (Supplementary Fig. S3) in the Z *tritici* treatment group (barcode 2) which displayed the most *Parastagonospora nodorum* assigned sequencing reads (Fig. 4). The *Puccinia striiformis* f. sp. *tritici* treatment group (barcode 1) and the *Puccinia striiformis* f. sp. *tritici + Z. tritici* treatment group (barcode 4) contained a lesser number of *Parastagonospora nodorum-*assigned sequencing reads, and these appeared to be below the detection threshold of PCR analysis. These results support the finding that *Parastagonospora nodorum* is present in our samples from the Wagga Wagga wheat-growing region despite the disease not being commonly found in the region nor the pathogen being purposefully inoculated in the trial.

**Fig. 3.**
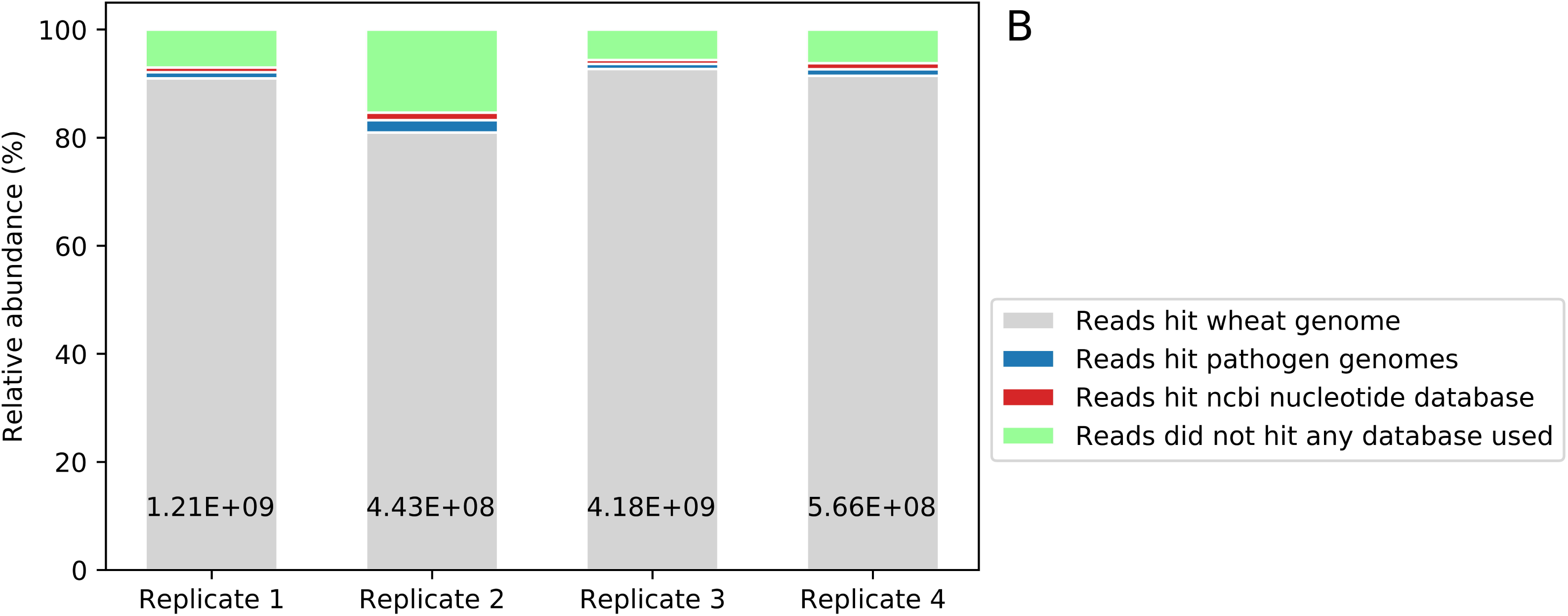
Wheat is predominantly identified in the whole genome sequencing analysis using BLAST. The graph shows the relative sequence length per species identified using a ‘winner-takes-all’ approach when searching the custom genome reference database. Wheat (grey), fungal pathogen genomes (blue), NCBI nucleotide (nt) database (red) and unmapped (light green) (see results and methods for details). The numbers at the bottom of each bar show the total sequence length (bp) of processed reads for each replicate. Replicates 1, 3, 4 showed similar classification levels while replicate 2 contained a higher number of unclassified sequences.

**Fig. 4.**
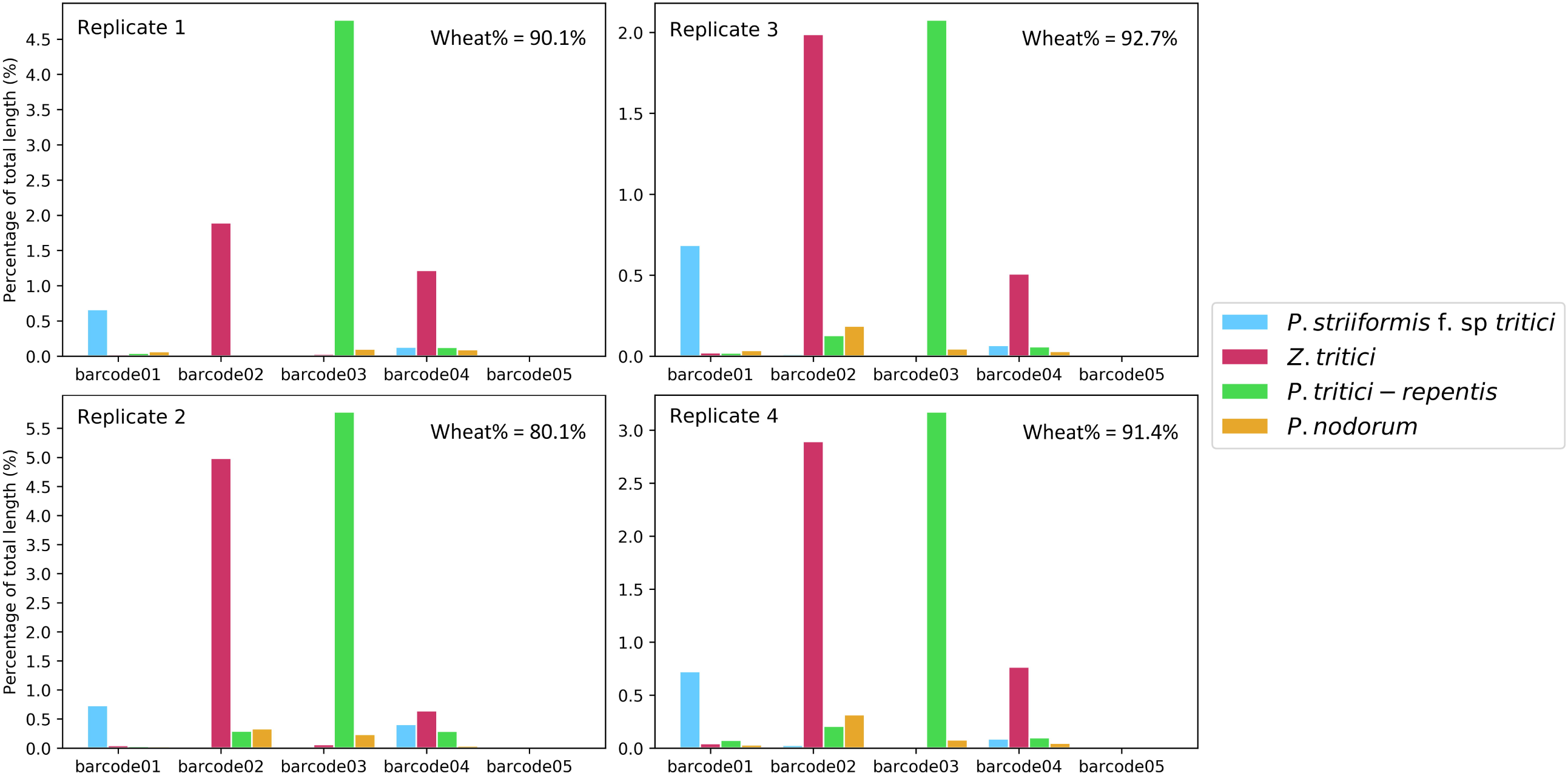
The total sequence length per pathogen genome per barcode correlated with the phenotypic disease identification. The graphs show the relative sequence length assigned to each pathogen genome using a ‘winner-takes-all’ approach in the BLAST analysis against the custom genome reference database. The proportion of reads hitting the wheat host genome are listed in the top-right corner of each graph. Unassigned reads are omitted from the visualization.

### Bacterial species associated with fungal disease development

To obtain an overview of the microbiome profile in and on wheat leaves infected by different fungal pathogens, we attempted to identify other microbial species present on the leaves. We searched the non-redundant NCBI nt database with reads that were not assigned to any of the five reference genomes present in our initial custom genome reference database. We found that the proportion of reads assigned to a genome in the restricted and the nt-NCBI databases were high for flowcells 1, 3 and 4 (92.9, 94.3 and 93.8%, respectively), but only 84.6% for flowcell 2 likely due to technical issues. Overall, from each flowcell, less than 1.4% of passed reads identified microbiota other than the major pathogens in the custom genome reference database. We pooled the reads from each barcode in all biological replicates, and calculated the total sequence length percentage that identified a specific genus in each sample using NCBI taxonomic identifiers (Fig. 5A). Genus *Pseudomonas* took up the highest proportion in all diseased samples, ranging from 0.08% to more than 0.88%, but was nearly absent in healthy wheat (barcode 5, 0.01%). Genus *Alternaria* is also found in all samples with a proportion between 0.05% to 0.2% in diseased samples and 0.005% in healthy wheat. Uniquely in *Puccinia striiformis* f. sp. *tritici* treatment group (barcode 1), bacterial *Erwinia* comprised 0.09% of total sequence length as represented by 219 reads and the second highest proportion. In contrast, *Erwinia* only represented 0.003% and 0.001% in Z *tritici* treatment group (barcode 2) and fungicide treatment group (barcode 5) while being absent in any other treatment group. In treatment groups infected with Z *tritici* (barcodes 2 and 4), hits on Z *tritici* were still present and may represent sequences in the six Z *tritici* genomes in the NCBI nt database that are absent from the Z *tritici* genome we used to generated our initial custom database. At the species level (Fig. 5B), total sequence length percentages identifying *Pseudomonas syringae* were much more frequent in treatment groups infected with *Puccinia striiformis* f. sp. *tritici* (barcode 1, 0.59%) and/or Z *tritici* (barcode 2, 0.33%). In contrast, the treatment group infected wtih *Pyrenophora tritici-repentis* (barcode 3) displayed a much lower relative total sequence length of 0.02% assigned to *Pseudomonas syringae.* Relative sequence length assigned to other *Pseudomonas* species such as *Pseudomonas poae* followed a similar trend. This finding is consisted with previous reports were *Pseudomonas* spp. have been descripted to be associated with fungal infections of wheat (Al-Sallami et al. 1997; Mehrabi et al. 2016). At the upper taxonomic classifications, *Proteobacteria* and *Ascomycota* were the most abundant bacterial and fungal phyla across all treatment groups (Supplementary Fig. S4) followed by *Actinobacteria* and *Basidiomycota.* At the class level (Supplementary Fig. S5), either *Gammaproteobacteria* or *Dothideomycetes* were the most abundant. Overall, the microbiome structure of the upper taxonomy resembles that of the lower taxonomy (Supplementary Fig. S4, Fig. S5, Fig. S6, Fig. S7), as most sequences were assigned to genus *Pseudomonas.*

**Fig. 5.**
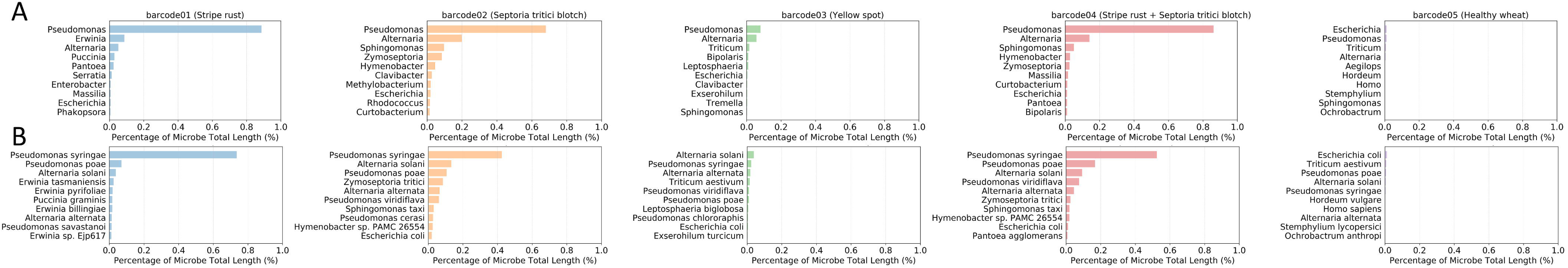
Fungal infections of wheat increase the microbial population, especially for *Puccinia striiformis* f. sp. *tritici* and Z *tritici* infections. Percentage of sequences that did not hit the custom genome reference database but hit the non-redundant NCBI nt database combined from all replicates, and grouped by genus (A) or species (B). Different colours represented different treatment types. Reads that were assigned to the restricted database were eliminated, and the top ten highest classifications were displayed on the y-axis for each replicate. Genus *Pseudomonas* represented the highest proportion across all barcodes containing *Puccinia striiformis* f. sp. *tritici* or Z *tritici,* with *Pseudomonas syringae* as the highest at the species level.

## DISCUSSION

We have developed a whole genome shot-gun sequencing (WGS) workflow which accurately identified three fungal pathogens responsible for three major wheat diseases - stripe rust, septoria tritici blotch and yellow spot - from field samples. We designed a two-step BLAST process to identify major agents in diseased samples and characterise the associated microbiota. We identified a number of *Pseudomonas* species present in stripe rust and septoria tritici blotch infected wheat leaves which appear far less abundant in yellow spot or non-diseased leaves.

WGS provides a wealth of information that could be used to accurately identify disease-causing agents and begin to describe microbial community composition in a sample (Brown et al. 2017; Thomas, Gilbert, and Meyer 2012). Without PCR amplification process to potentially introduce bias, WGS data is obtained directly from sample extracts and illustrates features closest to reality dependent only on sampling and DNA extraction strategies. Omitting PCR steps, WGS requires less processing time and proves advantageous for on-site diagnostic purposes where technical simplicity and fast processing time are prioritised. It has a significant drawback, however, in that most sequences are derived from the host (wheat) genome, representing 90.1% of total sequence length in flowcell 1, 80.1% in flowcell 2, 92.7% in flowcell 3 and 91.4% in flowcell 4. This result is due to our DNA extraction method in which the whole leaf sample was ground to retain all the microbial population in and on the leaf. For the purpose of microbial species identification, host sequences are unwanted and can overwhelm community DNA (Sharpton 2014). To enrich microbial sequences, one strategy is to apply intricate molecular methods during DNA extraction such as gradient centrifugation and sonication (Bulgarelli et al. 2012; Lim et al. 2014). As these techniques require sophisticated instruments, this can be a challenge to apply to environmental samples, especially in field conditions. Another strategy to enrich microbiome sequences involves sequencing of microbial metabarcode markers. With the improvement of sequencing length, species-level resolution using long read amplicon sequencing has been achieved in few cases (Benítez-Páez and Sanz 2017; Martijn et al. 2017). Long-read amplicon sequencing seems to be a promising method for species identification of known microbes, as it requires less input DNA than WGS and metabarcode databases contain information from comparatively more species than whole genome databases (Cruaud et al. 2017; Tessler et al. 2017; Ranjan et al. 2016). Amplicon sequencing is also possible in applied cases given the rapid development of remote PCR such as the miniPCR system (Amplyus, Cambridge, USA) and the Bento Lab platform (Bento Bioworks Ltd, London, United Kingdom). Drawbacks of amplicon strategy include biases related to PCR amplification, which lead to inaccuracy when calculating relative proportions of microbial species.

While many workflows are available for microbiome profiling using metagenomics data, there is no standard method for analysing long read data. This makes validation of both methods and results crucial (Quince et al. 2017), and mock microbial communities or samples containing known microbial species have been used to validate methods as a proof-of-concept (Nicholls et al. 2018). Verifying the diagnosis of a specific pathogen strain based on SNPs called from MinION reads, Votintseva et al. (2017) first identified pathogens using clinical methods and then generated sequencing data to map this data to the targeted species in real-time. Sequencing reactions were stopped once sufficient SNPs were obtained to confidentially identify the targeted pathogen strain. Similarly, our study used wheat samples inoculated manually by known pathogens and confirmed visually prior to sequencing. The major pathogen identifications from the first BLAST analysis against the custom genome reference database validated the capacity of our workflow for identification of those pathogen species. To validate the result, different analysis approaches should be applied on the same dataset. Brown et al. (2017) compared the performance of four bioinformatics pipelines on identifying species within different synthetic bacterial communities using MinION. Compared to the composition of a mock community, even the pipeline that resulted in the most similar classification still contains reads classified to species known not to be present. These false positive results are due largely to sequencing errors, which illustrates the need for more stringent filtering of the raw data and improvement of per read accuracy.

Nanopore metagenomics has great potential for quick diagnosis of suddenly emerging crop disease and large-scale disease monitoring at centralized agricultural institutions. As shown from other sequencing studies performed in a more extreme environment (Johnson et al. 2017; Boykin et al. 2018), all processes including DNA extraction and purification, library preparation and sequencing can be completed within a few hours. A tailored database can be created for each crop species containing the genomes of pathogens known to cause diseases to minimise processing time. For rapidly evolving pathogens such as wheat rusts, the SNP-based classification method described by Votintseva et al. (2017) could be adapted to define rust strains once enough SNPs are obtained. However, before running the sequencer, sampling procedures and tissue processing strategies still need to be carefully curated. In our study, we randomly sampled multiple leaves from different plants belonging to five treatment groups and performed complete sequencing and analysis independently on all four biological replicate sample sets. Consistency in barcode classification and BLAST analysis across all replicates suggested reliability of the workflow. In centralized agricultural institutions, amplicon sequencing can potentially play a key role and, to-date, there have been a few preliminary successes (Benítez-Páez, Portune, and Sanz 2016; Benítez-Páez and Sanz 2017; Pomerantz et al. 2018). Given the multiplexing capacity of the 96 native barcoding kit (ONT), many samples can be sequenced in parallel when larger scale sequencing platforms are employed (e.g. GridION, ONT). To achieve this goal, long read databases, sequencing accuracy and access to large-scale sample processing power need to be improved.

In conclusion, our workflow demonstrates the potential of this technology for plant pathogen diagnosis, field applications and microbiome characterisation. A combination of on-site and centralized sequencing approaches would, in future, revolutionize management of agricultural biosecurity and reduce crop loss.

## ACKNOWLEDGEMENTS

We thank Dr. Megan McDonald for providing photos of infected wheat leaves, and thank Mr. Michael McCaig, Mr. Tony Goldthorpe for their help during field sampling. We also thank Prof. Peter Solomon for sharing knowledge and PCR primers for *Parastagonospora nodorum Tox3* gene.

---

Supplementary Fig. S1. Example of DNA quality control by 1% agarose gel electrophoresis. About 200 nanograms of DNA from replicates 3 and 4 prior to the adapter ligation step of the sequencing library preparation were loaded onto the gel. The molecular weight standards are HyperLadder 1 kb (Bioline, Catalog number BIO-33025).

Supplementary Fig. S2. BLAST analysis against the non-redundant NCBI nt database using the reads assigned to *Parastagonospora nodorum* from BLAST against the restricted database. Arc angle is proportional to the number of reads assigned to the indicated genus or order. The unclassified (grey) arc angle refers to the proportion of reads assigned as unclassified by BLAST analysis. Genus or orders assigned to two reads or less are grouped as ‘others’ under each category.

Supplementary Fig. S3. PCR amplification of the *Parastagonospora nodorum Tox3* gene from the DNA sample of replicate 3. The wheat *Rubisco* gene was used as a control for the quality of DNA in each sample. The positive control for *Tox3* amplification is pure genomic DNA from *Parastagonospora nodorum* strain SN15, and the negative control is pure genomic DNA from Z *tritici.* The PCR reaction used NEB Q5 High-Fidelity 2X Master Mix (Catalog Number M0492S) following the manufacturer’s protocol and the amplification conditions were: 30 s at 98°C; 30 cycles of 10 s at 98°C, 25 s at 56°C and 10 s at 72°C; then a final extension for 2 min at 72°C before holding at 4°C. The primer sequences for the wheat *Rubisco* gene are: Forward: AGCAAGGTTGGCTTTGTCTT, Reverse: TTAACCTCCTCCACCTCGTT. The primer pair used for the *Tox3* gene are: Forward: A AT G TCG ACCG TTTT G ACC, Reverse: GGTTGCCGCAGTTGATATAA obtained from Lin, Chooi, and Solomon (2018).

Supplementary Fig. S4. Percentage of sequences that did not hit the custom genome reference database but hit the non-redundant NCBI nt database combined from all replicates, and grouped by phylum. Reads that are assigned to the restricted database were eliminated, and the top ten highest classifications were displayed for each replicate. Different colours represented different barcodes.

Supplementary Fig. S5. Percentage of sequences that did not hit the custom genome reference database but hit the non-redundant NCBI nt database combined from all replicates, and grouped by class. Reads that are assigned to the restricted database were eliminated, and the top ten highest classifications were displayed for each replicate. Different colours represented different barcodes.

Supplementary Fig. S6. Percentage of sequences that did not hit the custom genome reference database but hit the non-redundant NCBI nt database combined from all replicates, and grouped by order. Reads that are assigned to the restricted database were eliminated, and the top ten highest classifications were displayed for each replicate. Different colours represented different barcodes.

Supplementary Fig. S7. Percentage of sequences that did not hit the custom genome reference database but hit the non-redundant NCBI nt database combined from all replicates, and grouped by family. Reads that are assigned to the restricted database were eliminated, and the top ten highest classifications were displayed for each replicate. Different colours represented different barcodes.

