## Supplementary Table S1 for "Pathogen Detection and Microbiome Analysis of Infected Wheat Using a Portable DNA Sequencer"

Supplementary Table S1. Basic statistical comparison of each sequencing replicate. The barcode column indicates DNA barcodes used in the library preparation for each replicate.

| **Replicate** | **No. of Q^a^ > 7 reads/ No. of total reads (bp^b^, %)** | **Q > 7 total read length/ Total read length (bp, %)** | **Total read length after Porechop (bp)** | **Mean read length from Q > 7 reads (bp)** | **Barcodes** | **Disease confirmed by pathologist** |
| --- | --- | --- | --- | --- | --- | --- |
| 1 | 847,469/983,698 (86.2%) | 1,587,046,441/1,707,799,548 (92.9%) | 1,206,433,874 | 1872.7 | barcode02 | stripe rust |
|  |  |  |  |  | barcode03 | yellow leaf spot |
|  |  |  |  |  | barcode04 | None |
|  |  |  |  |  | barcode05 | stripe rust and septoria tritici blotch |
|  |  |  |  |  | barcode06 | septoria tritici blotch |
| 2 | 568,686/659,039 (86.3%) | 785,073,330/837,189,225 (93.8%) | 442,667,426 | 1380.5 | barcode07 | stripe rust |
|  |  |  |  |  | barcode08 | septoria tritici blotch |
|  |  |  |  |  | barcode09 | None |
|  |  |  |  |  | barcode10 | stripe rust and septoria tritici blotch |
|  |  |  |  |  | barcode11 | yellow leaf spot |
| 3 | 1,932,416/2,123,611 (91.0%) | 5,051,730,065/5,304,946,873 (95.2%) | 4,175,446,769 | 2614.2 | barcode01 | stripe rust |
|  |  |  |  |  | barcode02 | septoria tritici blotch |
|  |  |  |  |  | barcode03 | None |
|  |  |  |  |  | barcode04 | stripe rust and septoria tritici blotch |
|  |  |  |  |  | barcode05 | yellow leaf spot |
| 4 | 223,192/242,106 (92.2%) | 681,656,642/736,410,201 (92.6%) | 565,771,763 | 3054.1 | barcode01 | stripe rust |
|  |  |  |  |  | barcode02 | septoria tritici blotch |
|  |  |  |  |  | barcode03 | None |
|  |  |  |  |  | barcode04 | stripe rust and septoria tritici blotch |
|  |  |  |  |  | barcode05 | yellow leaf spot |

^a^ Q: Phred quality

^b^ bp: basepair
