## Supplementary figures and images for "Pathogen Detection and Microbiome Analysis of Infected Wheat Using a Portable DNA Sequencer"

### Fig. S5

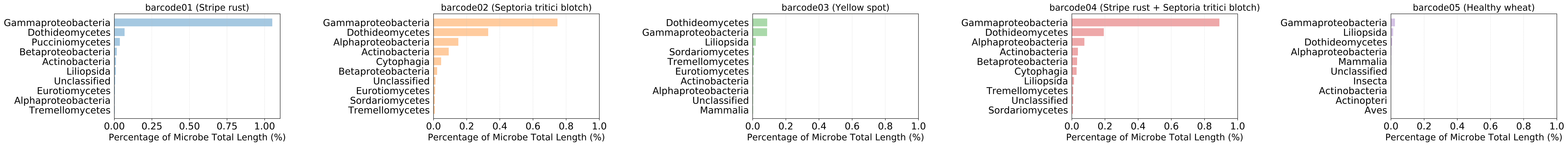

### Fig. S6

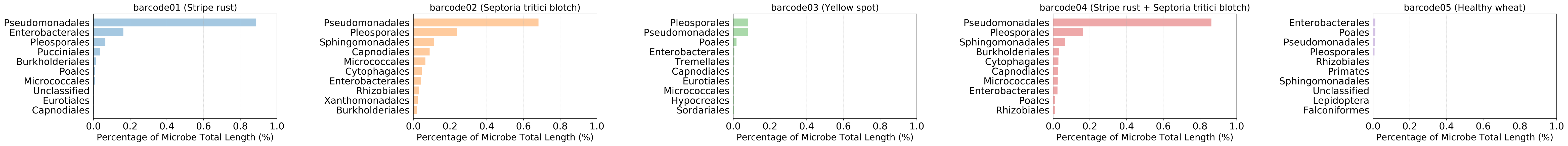

### Fig. S7

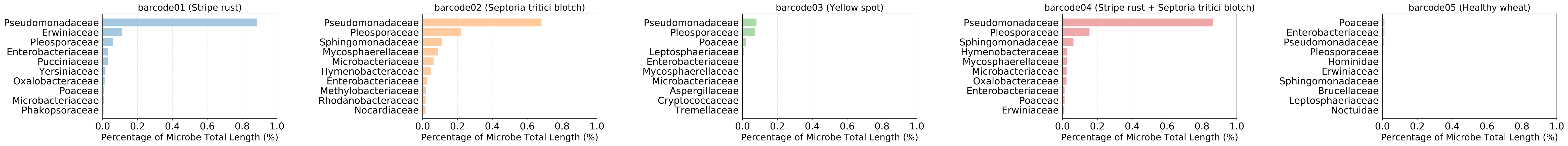

### Supplementary Fig. S1

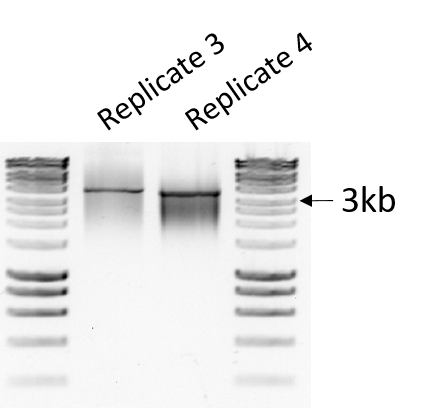

### Supplementary Fig. S2

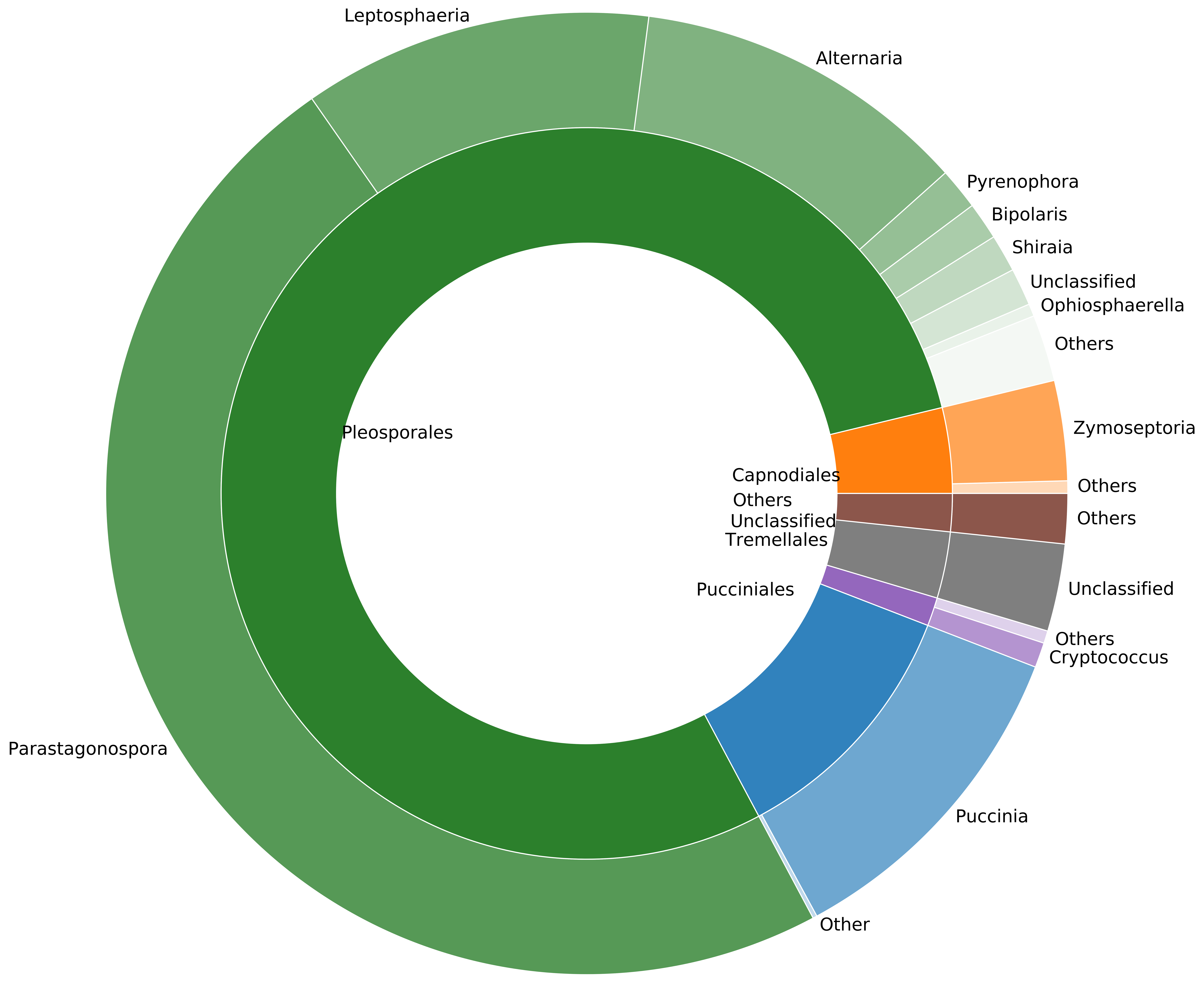

### Supplementary Fig. S3

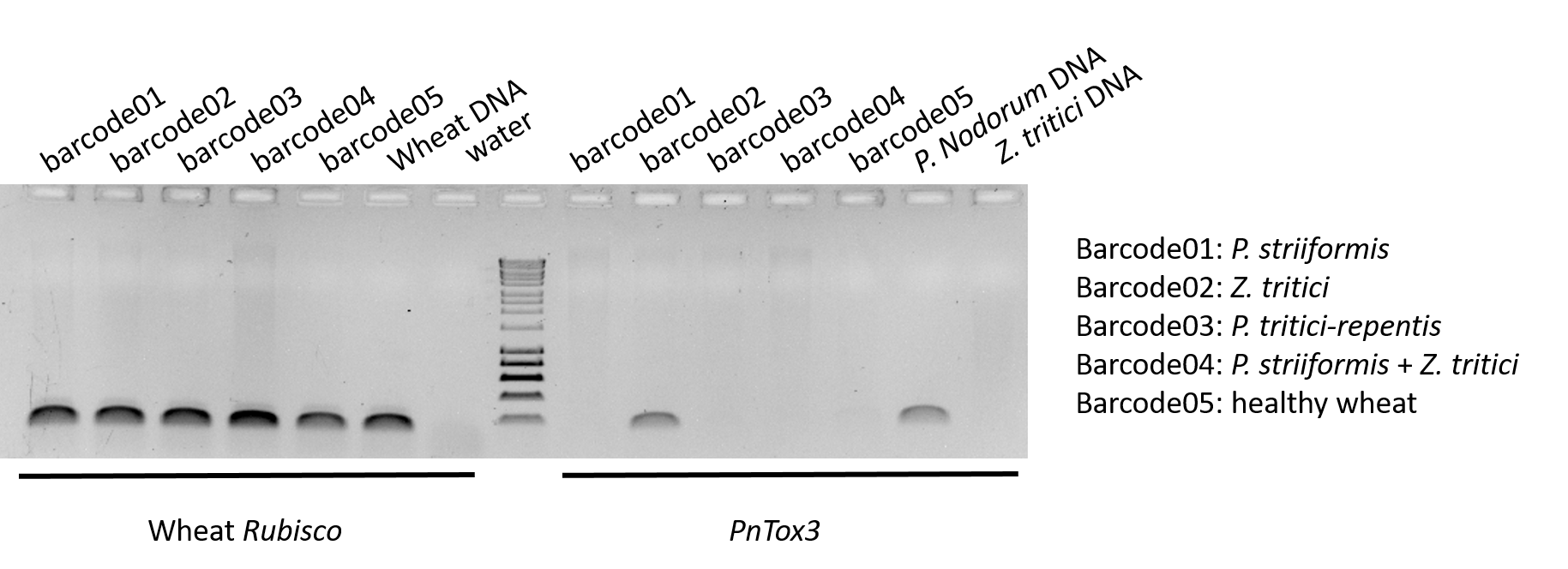

### Supplementary Fig. S4

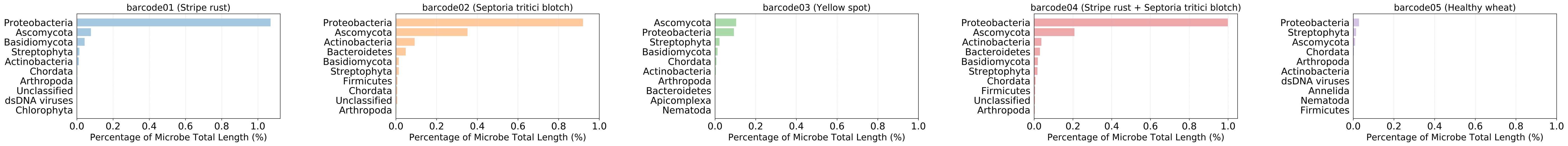
